# Computational correction of copy-number effect improves specificity of CRISPR-Cas9 essentiality screens in cancer cells

**DOI:** 10.1101/160861

**Authors:** Robin M. Meyers, Jordan G. Bryan, James M. McFarland, Barbara A. Weir, Ann E. Sizemore, Han Xu, Neekesh V. Dharia, Phillip G. Montgomery, Glenn S. Cowley, Sasha Pantel, Amy Goodale, Yenarae Lee, Levi D. Ali, Guozhi Jiang, Rakela Lubonja, William F. Harrington, Matthew Strickland, Ting Wu, Derek C. Hawes, Victor A. Zhivich, Meghan R. Wyatt, Zohra Kalani, Jaime J. Chang, Michael Okamoto, Todd R. Golub, Jesse S. Boehm, Francisca Vazquez, David E. Root, William C. Hahn, Aviad Tsherniak

## Abstract

The CRISPR-Cas9 system has revolutionized gene editing both on single genes and in multiplexed loss-of-function screens, enabling precise genome-scale identification of genes essential to proliferation and survival of cancer cells. However, previous studies reported that an anti-proliferative effect of Cas9-mediated DNA cleavage confounds such measurement of genetic dependency, particularly in the setting of copy number gain^1-4^. We performed genome-scale CRISPR-Cas9 essentiality screens on 342 cancer cell lines and found that this effect is common to all lines, leading to false positive results when targeting genes in copy number amplified regions. We developed CERES, a computational method to estimate gene dependency levels from CRISPR-Cas9 essentiality screens while accounting for the copy-number-specific effect, as well as variable sgRNA activity. We applied CERES to sets of screens performed with different sgRNA libraries and found that it reduces false positive results and provides meaningful estimates of sgRNA activity. As a result, the application of CERES improves confidence in the interpretation of genetic dependency data from CRISPR-Cas9 essentiality screens of cancer cell lines.

---

Major efforts using loss-of-function genetic screens to systematically identify genes essential to the proliferation and survival of cancer cells have been reported^1-9^. Genes identified by these approaches may represent specific genetic vulnerabilities of cancer cells, suggesting treatment strategies and directing the development of novel therapeutics. The CRISPR-Cas9 genome editing system has proven to be a powerful tool to interrogate gene essentiality in cancer cell lines. Its relative ease of application, high rates of target validation, and increased specificity compared to RNA interference technology make it an ideal instrument for use in high-throughput functional genomic screening^10^.

However, we and others have recently observed that measurements of genetic dependency in genome-scale CRISPR-Cas9 loss-of-function screens are influenced by the genomic copy number (CN) of the region targeted by the sgRNA-Cas9 complex^1-4^. Targeting Cas9 to DNA sequences within regions of high CN gain creates multiple DNA double-strand breaks (DSBs), inducing a gene-independent DNA damage response and a G2 cell-cycle arrest phenotype^2^. This systematic, sequence-independent effect due to DNA cleavage (*copy-number effect*) confounds the measurement of the consequences of gene deletion on cell viability (*gene-knockout effect*) and is detectable even among low-level CN amplifications and deletions. In particular, this phenomenon hinders interpretation of CRISPR-Cas9 experiments in cancer cell lines, which are frequently aneuploid and harbor large numbers of genomic amplifications and deletions. In these cell lines, amplified genes represent a major source of false positives^2,3^. Existing methods to handle the copy-number effect adopt filtering schemes^4^, which preclude examination of data from amplified regions and ignore the copy-number effect at low level alterations. Here, we present CERES – a method to estimate gene dependency from essentiality screens while computationally correcting the copy-number effect – enabling unbiased interpretation of gene dependency at all levels of CN.

As part of our efforts to build a Cancer Dependency Map, a catalog of cell line-specific genetic and chemical vulnerabilities, we performed genome-scale CRISPR-Cas9 loss-of-function screens in 342 cancer cell lines representing 27 cell lineages **(Supplementary Table 1**) using the Avana sgRNA library^11^ (**Supplementary Table 2**) and assessed the effects of introducing each sgRNA on cell proliferation (Online Methods). After applying quality control measures, ROC analysis of sgRNAs targeting common core essential and nonessential genes^12^ demonstrated high screen quality in all cell lines (**Fig. 1a**). We also reanalyzed published datasets of 33 cancer cell lines of diverse cell lineage (GeCKOv2) ^2^ and 14 AML cell lines (Wang2017) ^4^ (**Supplementary Fig. 1a**).

**Figure 1:**
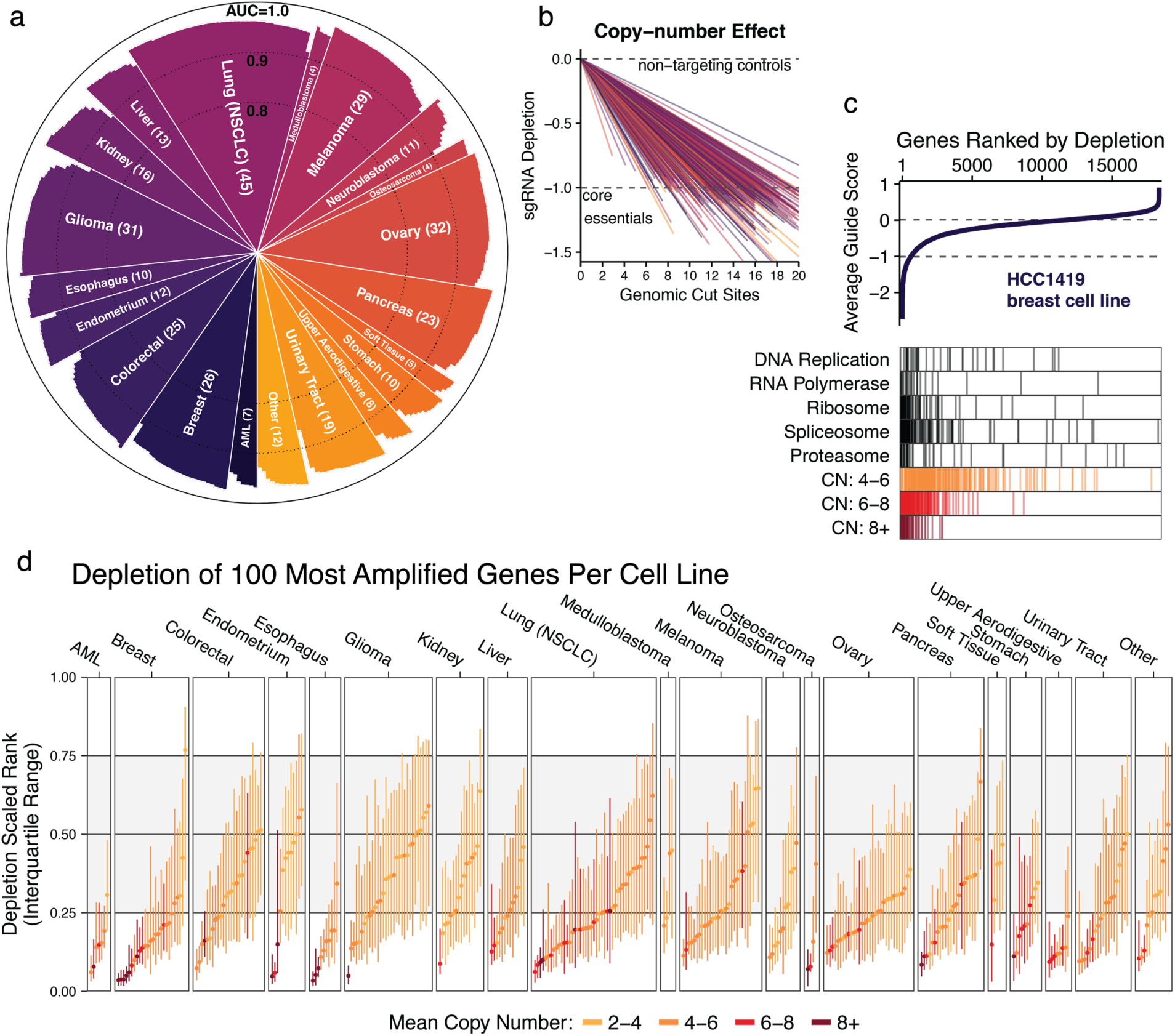
Genomic copy number confounds the interpretation of CRISPR-Cas9 loss-of-function proliferation screens of cancer cell lines. (**a**) Screen quality for each cell line in the panel (n = 342), as measured by area under the receiver operating characteristic curve (AUC) in discriminating between predefined sets of common core essential and nonessential genes. (**b**) For each cell line, the depletion of sgRNAs is regressed against the number of perfect-match genomic cut sites using a piecewise-linear fit. The slope of the fit is plotted and represents the average effect per expected cut on cell proliferation. Each cell line is scaled such that the median of sgRNAs targeting cell-essential genes is at −1, marked by a dashed line. (**c**) Genes are ranked by the mean depletion of targeting sgRNAs (average guide score) and plotted for an example cell line. A value of −1 represents the median of cell-essential genes, indicated by a dashed line. Below, depletion ranks of genes involved in fundamental cell processes and genes at various ranges of CN amplification are shown. (**d**) The median and interquartile range (IQR) of depletion ranks for the 100 most amplified genes per cell line are plotted. Color indicates mean amplification level of these genes. The grey shaded area indicates the IQR of all genes.

The copy-number effect was characterized in previous efforts in a limited number of cell contexts with measurements using different sgRNA libraries. We assessed the 342 cell lines screened in our dataset for sensitivity to Cas9-mediated cleavage as in Aguirre *et al.* ^2^. In consonance with previous observations, every cell line in our panel was sensitive to the copy-number effect, where sgRNAs targeting more genomic loci were on average more depleted, frequently to levels at or below the depletion of sgRNAs targeting cell-essential genes (**Fig. 1b, Supplementary Fig. 1b**). While this relationship held in all cell lines, some variability in the strength of effect could be explained by *p53* mutational status (**Supplementary Fig. 1c**).

To determine how this sgRNA-level effect translates into false positive gene dependencies, we ranked the genes in each cell line by the average depletion of their targeting sgRNAs (*average guide score*). In an example breast cancer cell line, HCC1419, high-ranking genes were enriched for both genes involved in fundamental cellular processes and genes with amplified CN (**Fig. 1c**). The depletion ranks of the 100 genes with the largest CN measurements were significantly higher than expected for the majority of cell lines (300/342 with FDR-corrected p < 0.05, one-sample one-tailed K-S test; **Fig. 1d**) and the extent of enrichment was significantly correlated with the average CN of these genes (Spearman ρ = 0.60, p < 2.2 x 10^-16^, **Supplementary Fig. 2a**), consistent with previous studies (**Supplementary Fig. 2b**).

In order to decouple gene dependency from the effects of Cas9-mediated cleavage in sgRNA depletion data, CERES models each measured depletion value as a sum of unknown gene-knockout and copy-number effects (**Fig. 2**). The copy-number effect is a function of the number of DNA cuts induced by the sgRNA, accounting for potential multiple alignments to the genome and the CN at each locus. The sum of these two effects is multiplied by a *guide activity score*, which estimates the degree to which the sgRNA induces the expected depletion effects and was included to lessen the impact of low-quality reagents^11,13,14^. CERES infers the gene-knockout effects, which represent the underlying gene dependencies in each cell line, as well as the copy-number effects and the guide activity scores by fitting the above model to sgRNA depletion and CN data (Online Methods).

**Figure 2:**
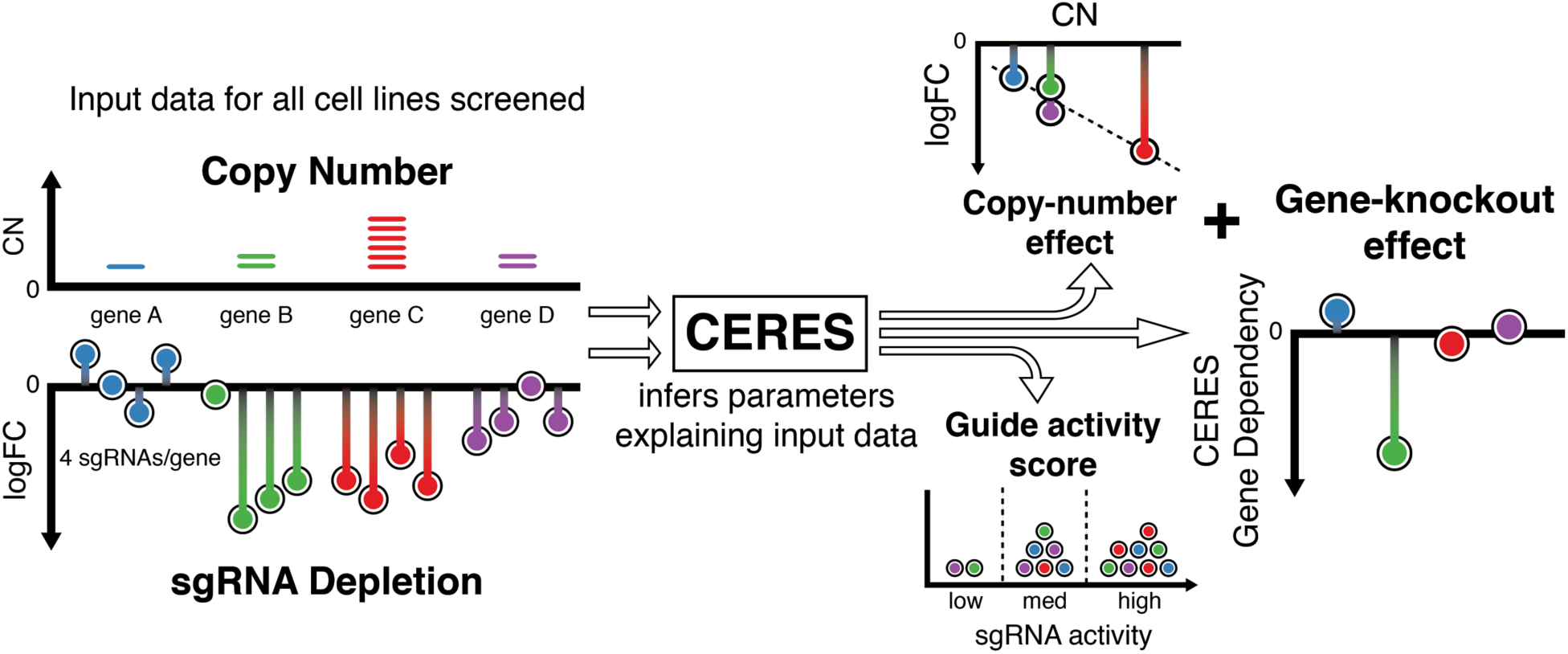
Schematic of the CERES computational model. As input, CERES takes sgRNA depletion and CN data for all cell lines screened. During the inference procedure, CERES models the depletion values as a sum of gene-knockout and copy-number effects, multiplied by a guide activity score parameter. CERES then outputs the values of the parameters that produce the highest likelihood of the observed data under the model.

We applied CERES to our dataset of 342 essentiality screens and assessed the performance of the model by comparing CERES gene dependency scores to the uncorrected average guide scores. As expected, CERES markedly reduced the relationship between CN and gene dependency found in the uncorrected average guide scores (**Fig. 3a, Supplementary Fig. 3a**). We correlated dependency scores for each gene to its CN measurements before and after correction and found that CERES shifted the mean correlation to near zero (**Supplementary Fig. 3b**). CERES also improved the identification of essential genes in all 342 screens, as measured by the recall of cell-essential genes at a 5% false discovery rate (FDR) ^8^, by an average of 21.8 percentage points (**Fig. 3b, Supplementary Fig. 4**) (Online Methods). Reassuringly, CERES preserved expected cancer-specific dependencies, even in amplified regions. In an example *KRAS* amplification on chromosome 12p of the DAN-G pancreatic cancer cell line, CERES removed the local enrichment of gene dependencies while preserving the essentiality of *KRAS* (**Fig. 3c**, **Supplementary Fig. 5**). Additionally, *KRAS*-mutant cell lines remained substantially enriched over wild type for *KRAS* gene dependency (**Fig. 3d, Supplementary Fig. 6**).

**Figure 3:**
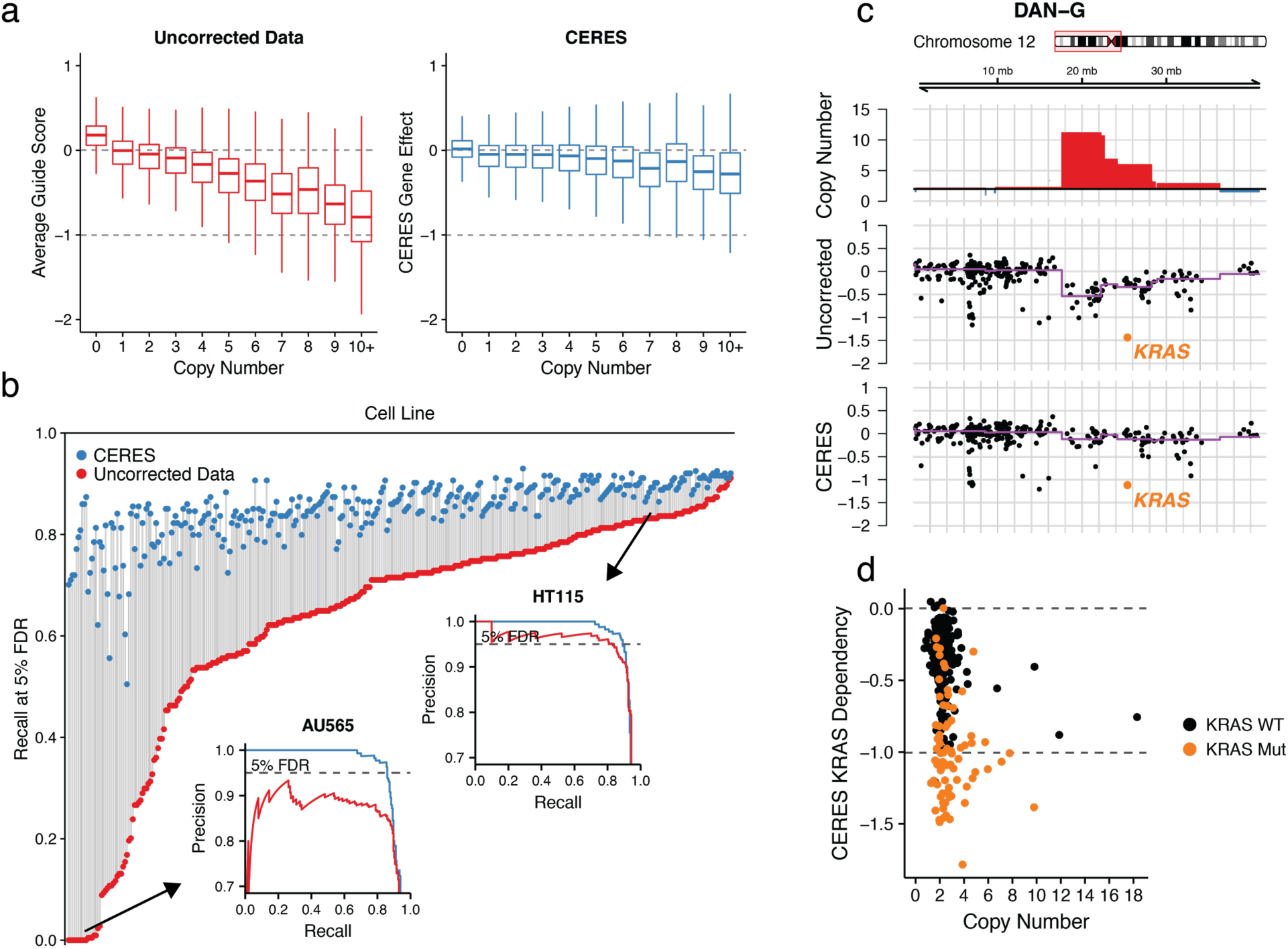
CERES corrects the copy-number effect and improves specificity of CRISPR-Cas9 essentiality screens, while preserving true gene dependencies. (**a**) Boxplots of gene dependency scores are shown across CN for uncorrected average guide scores and CERES gene dependency scores. Data are scaled as in Fig. 1b such that-1 represents the median score of cell-essential genes. (**b**) The recall of cell-essential genes at a 5% FDR of nonessential genes is plotted for each cell line before (red) and after (blue) CERES correction. Precision-recall curves are inset for example cell lines with poor recall (bottom left) and good recall (top right) before CERES correction. (**c**) An example amplified region on chromosome 12p is shown for the DAN-G pancreatic cell line. The top track represents CN with amplifications shown in red. The middle track shows the uncorrected average guide score for each gene in this region, with the purple trend line representing the median value in each CN segment. *KRAS* dependency is highlighted in orange. The bottom track shows the CERES gene dependency score, with the trend line as above. (**d**) *KRAS* gene dependency and CN are shown for all cell lines after CERES correction, with mutant *KRAS* lines in orange.

While it is infeasible to experimentally validate the activity of all sgRNAs in a genome-scale library, sequence determinants have proven useful in the prediction of on-target activity^11,15,16^. The Avana sgRNA library was optimized using such predictions. Fittingly, CERES estimated higher guide activity scores on average for the Avana dataset relative to GeCKOv2, with a near nine-fold increase in the ratio of high-to low-activity sgRNAs (**Fig. 4a**). The guide activity scores for the 5,044 sgRNAs shared between the two libraries showed substantial agreement (Spearman ρ = 0.53, p < 2.2 x 10^-16^), demonstrating that CERES captured a measure of sgRNA activity that is reproducible across independent screens of cell line panels (**Fig. 4b, Supplementary Fig. 7**). For both the GeCKOv2 and Avana libraries, we compared CERES guide activity scores to sequence-based predictions of sgRNA activity (Doench-Root scores) and found significant correspondence (Avana: Pearson ρ = 0.22, p < 2.2 x 10^-16^; GeCKOv2: Pearson ρ = 0.44, p < 2.2 x 10^-16^; **Fig. 4c**). Taken together, this evidence demonstrates that the guide activity scores inferred by CERES are useful for estimating gene-knockout effects and, furthermore, suggests that these scores could be helpful in the selection of reagents for follow-up experiments.

**Figure 4:**
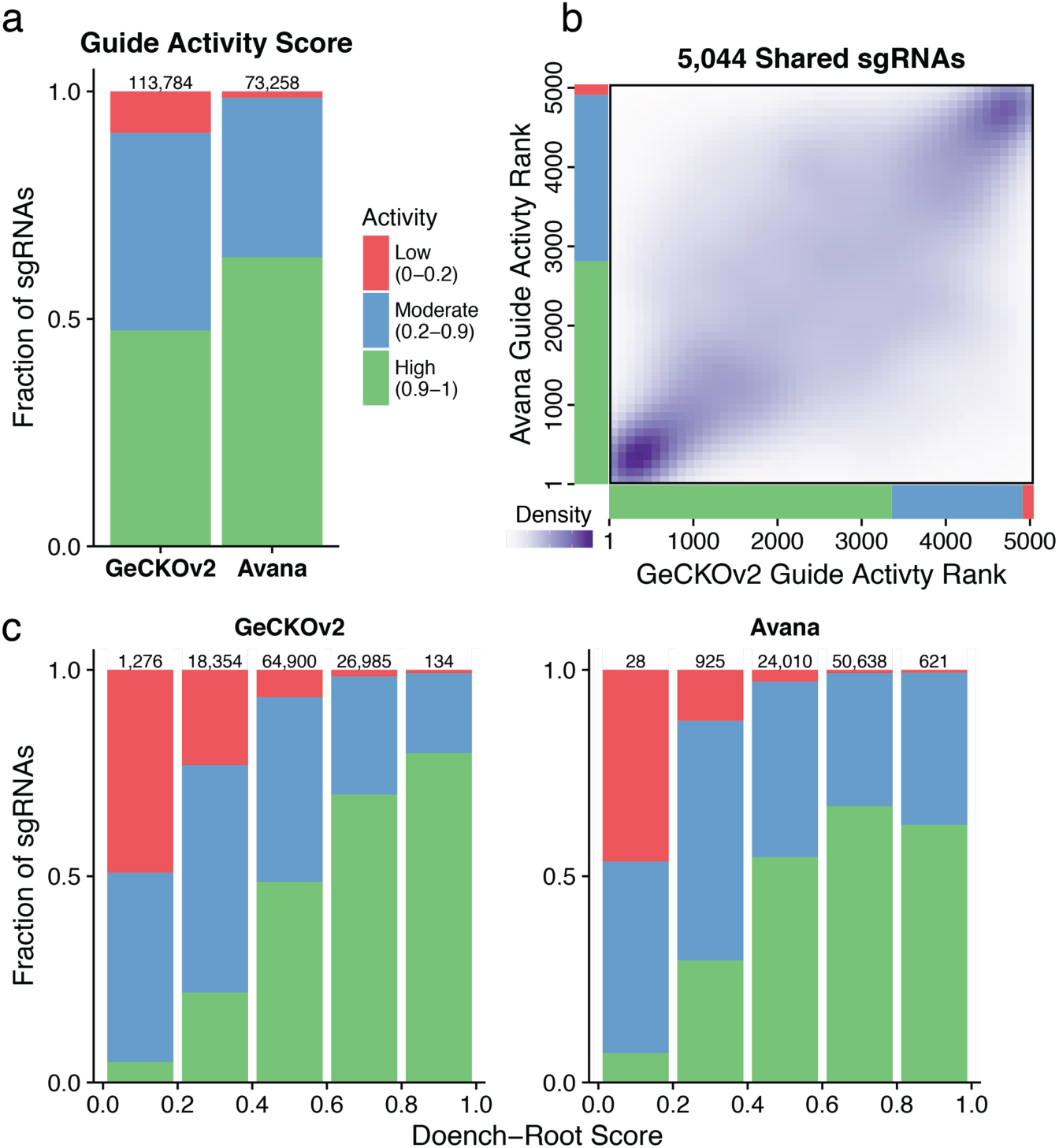
CERES infers guide activity scores for each sgRNA. (**a**) sgRNAs are binned into groups with high (0.9-1), moderate (0.2-0.9), and low (0-0.2) estimated activity scores. The compositions of guide activity scores are shown for the set of screens performed with the GeCKOv2 sgRNA library and the Avana sgRNA library, which is more optimized for on-target activity. (**b**) For the set of 5,044 sgRNAs shared between the GeCKOv2 and Avana libraries, sgRNAs are ranked by guide activity scores in each dataset and are plotted against each other, with darker purple representing greater density of sgRNAs. (**c**) sgRNAs are binned by predicted on-target activity using the Doench-Root score. For each dataset, the composition of CERES-estimated guide activity scores is shown for each Doench-Root bin.

To identify cancer-specific genetic vulnerabilities, we used a metric of differential dependency to represent the strength of dependency in a cell line relative to all other lines screened (Online Methods). We assessed an upper bound on the number of false positive differential dependencies due to CN amplifications by calculating the fraction of amplified genes at every threshold of differential dependency across our dataset. In the uncorrected data, the fraction of amplified genes increased at stronger dependency thresholds, climbing above 30% at the highest levels of differential dependency (**Fig. 5a, Supplementary Fig. 8a**). By contrast, CERES results maintained a low prevalence of amplified genes at every level of differential dependency. We next used a similar procedure to examine unexpressed genes, expected to be functionally inconsequential, which represent an overt source of false positives if scored as differentially dependent. We found that CERES reduced the fraction of unexpressed genes at high levels of dependency from 7% to 1%, indicating a substantial improvement in specificity (**Fig. 5b, Supplementary Fig. 8b**).

**Figure 5:**
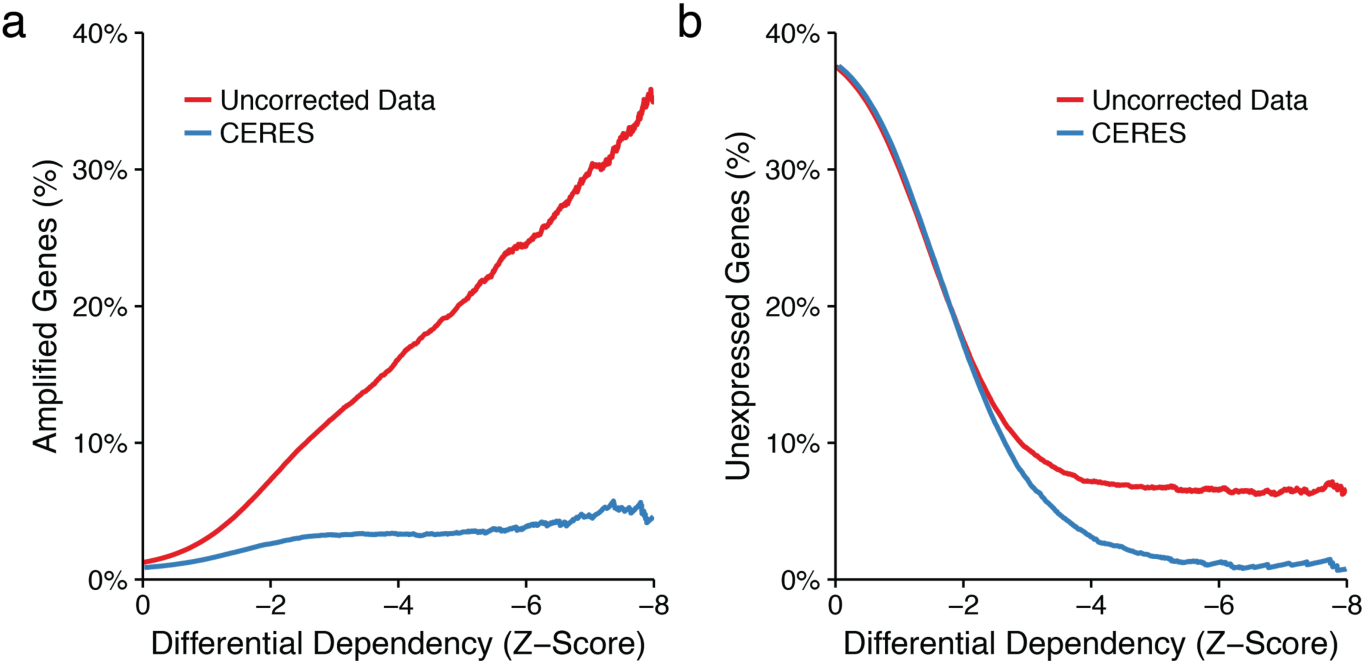
CERES reduces false positive differential dependencies. (**a**) For all cell lines in our dataset, the percentage of genes on amplified regions (CN > 4) below a given differential dependency threshold is plotted for the uncorrected average guide score in red and the CERES gene dependency score in blue. (**b**) The percentage of unexpressed genes (log_2_RPKM <-1) below a given differential dependency score is plotted as in (a).

A dataset of this scale enables the discovery of genetic vulnerabilities specific to a subset of cancer cell lines defined by some cellular context, such as cell lineage. We hypothesized that in this setting, copy-number effects driven by recurrent CN alterations, even with small effect sizes, could introduce false positives. For each gene, we compared average guide scores in breast cancer cell lines to those of all other cell lines (Online methods). Indeed, significant differential dependencies in breast lines were enriched on chromosome 8q, which is recurrently amplified in breast tumors (**Fig. 6a**). The same analysis applied to CERES-corrected dependency scores yielded two 8q genes with significant differential dependency: *TRPS1* and *GRHL2* (**Fig. 6b**). Previous studies have implicated these transcription factors in breast cancer progression^17,18^, and analysis of expression levels of these and other transcription factors suggest that they are likely to be truly differentially dependent in breast lines (**Supplementary Fig. 9**). We expanded this analysis to all cell lineages with recurrently amplified chromosome arms and quantified the enrichment of differential dependencies before and after CERES correction in each context. We observed that CERES reduced the fraction of differential dependencies on the recurrently amplified chromosome arm in 19 out of 24 such cases (**Fig. 6c**) (Online Methods).

**Figure 6:**
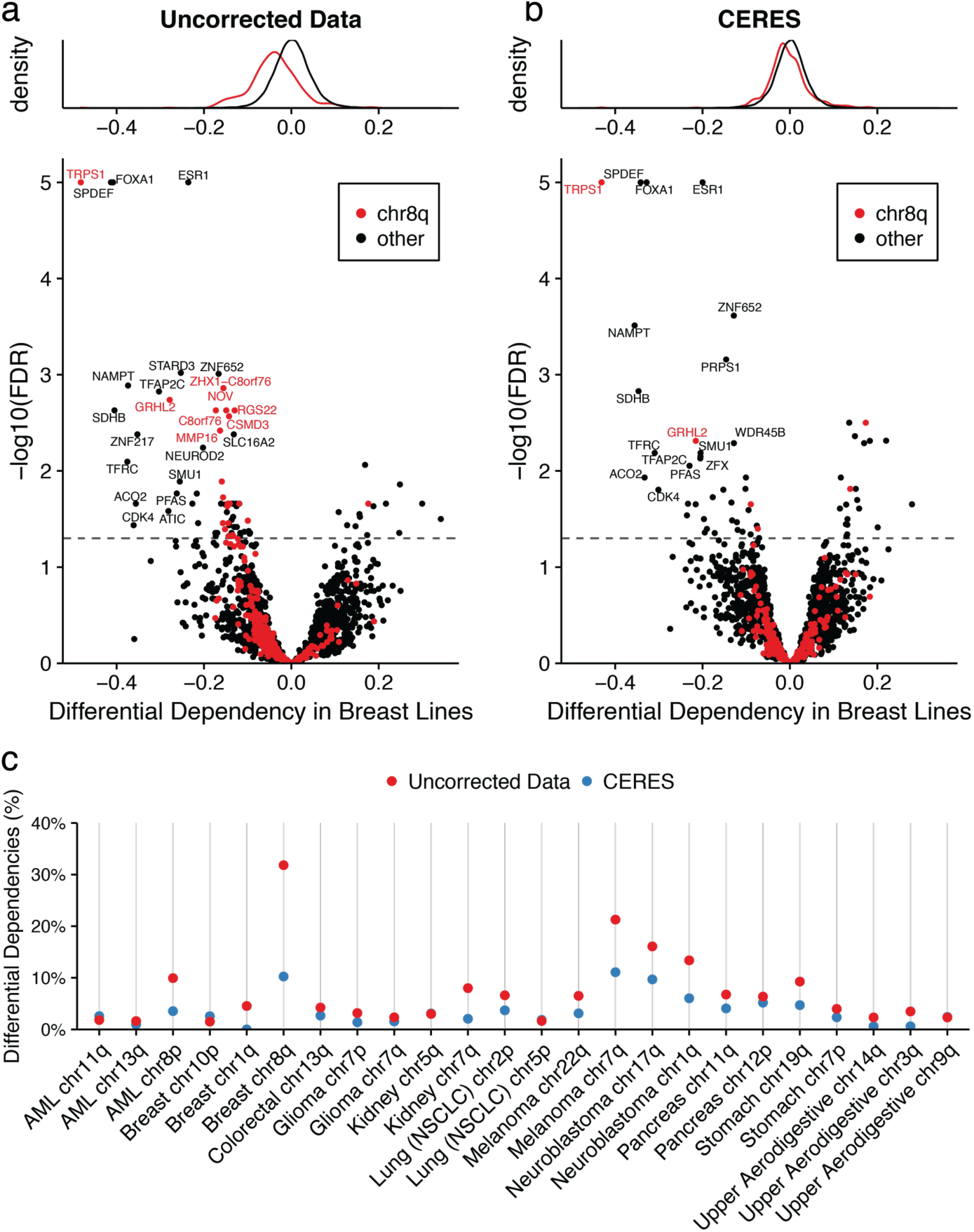
CERES reduces false positives in lineage-specific differential dependencies due to recurrently amplified chromosome arms. (**a**) Differential dependency in breast cancer cell lines is calculated as the difference in mean gene scores between breast lines and the rest of the cell lines screened. The distributions of differential dependencies in breast lines are plotted red for genes on chromosome 8q (commonly gained in breast tumors) and black for all other genes. Below, the differential dependency for each gene is plotted against the FDR-corrected p-value, calculated from a student’s t-test, with colors as above. The dashed line represents an FDR of 5%. (**b**) Data is plotted for CERES-inferred gene effects as in (a). (**c**) Percentage of lineage-specific differential dependencies (FDR < 0.05) that are on the specified chromosome arms is shown for arms that are recurrently amplified in those lineages, before and after CERES correction.

Here we introduce the largest set of CRISPR-Cas9 essentiality screens to date and propose a methodology to estimate gene dependency while correcting for copy-number effects. CERES removes false positive results due to the copy-number effect, revealing underlying genetic dependencies and enabling identification of essential genes with improved specificity.

## Acknowledgements

This project was supported by U01 CA176058, U01 CA199253, and P01 CA154303 grants (W.C.Hahn). This work was supported by the Slim Initiative for Genomic Medicine, a project funded by the Carlos Slim Foundation and the H.L. Snyder Foundation.

## Author contributions

R.M.M, J.G.B, and A.T. conceived of and designed the study. R.M.M., J.G.B., and J.M.M. performed computational analysis and interpretation of results. J.G.B. wrote and implemented the modeling software. R.M.M., B.A.W., and A.E.S. processed and managed data. H.X., and N.V.D. assisted with computational analysis. P.G.M. provided computational tools. G.S.C., S.P., and F.V. provided project management. A.G., Y.L., L.D.A., G.J., R.L., W.F.H., M.S., T.W., D.C.H., V.A.Z., M.R.W., Z.K., J.J.C., and M.O. assisted with data generation. R.M.M., J.G.B., J.M.M, W.C.H., and A.T. wrote and/or revised the manuscript with assistance from other authors. T.R.G., J.S.B., F.V., D.E.R., W.C.H., and A.T. supervised the study and performed an advisory role.

## Materials and correspondence

Correspondence or requests for materials should be addressed to either William C. Hahn or Aviad Tsherniak.

